# Functional or vestigial? The genomics of the pineal gland in Xenarthra

**DOI:** 10.1101/2021.05.17.444431

**Authors:** Raul Valente, Filipe Alves, Isabel Sousa Pinto, Raquel Ruivo, L. Filipe C. Castro

**Affiliations:** CIMAR/CIIMAR - Interdisciplinary Centre of Marine and Environmental Research, University of Porto, Avenida General Norton de Matos, S/N, 4450-208 Matosinhos, Portugal; FCUP - Department of Biology, Faculty of Sciences, University of Porto (U. Porto), Rua do Campo Alegre, Porto, Portugal; MARE - Marine and Environmental Sciences Centre, ARDITI, Madeira, Portugal; OOM - Oceanic Observatory of Madeira, Funchal, Portugal

**Keywords:** Gene Loss, Vestigiality, Pineal Gland, Xenarthra

## Abstract

Vestigial organs are historical echoes of past phenotypes. Determining whether a specific organ constitutes a functional or vestigial structure can be a challenging task, given that distinct levels of atrophy may arise between and within lineages. The mammalian pineal gland, an endocrine organ involved in melatonin biorhythmicity, represents a classic example, often yielding contradicting anatomical observations. In Xenarthra (sloths, anteaters and armadillos), a peculiar mammalian order, the presence of a distinct pineal organ was clearly observed in some species (i.e. Linnaeus’s two-toed sloth) but undetected in other closely related species (i.e. brown-throated sloth). In the nine-banded armadillo, contradicting evidence supports either functional or vestigial scenarios. Thus, to untangle the physiological status of the pineal gland in Xenarthra, we used a genomic approach to investigate the evolution of the gene hub responsible for melatonin synthesis and signaling. We show that both synthesis and signaling compartments are eroded and were lost independently. Additionally, by expanding our analysis to 157 mammal genomes we offer a comprehensive view showing that species with very distinctive habitats and lifestyles have convergently evolved a similar phenotype: Cetacea, Pholidota, Dermoptera, Sirenia and Xenarthra. Our findings suggest that the recurrent inactivation of melatonin genes correlates with pineal atrophy, and endorse the use of genomic analyses to ascertain the physiological status of suspected vestigial structures.

## 1. Introduction

Understanding the evolution of organ reduction, atrophy or indeed complete loss is a fascinating quest, dating back to the seminal work of Charles Darwin, *On the Origin of Species* (Darwin, 1859). Yet, to identify a structure as vestigial, described as a trait with no function, operating sub-optimally, or even with a modified function from that originally served, is no easy undertaking (Werth, 2014; Allmon & Ross, 2018): often yielding contradictory anatomical descriptions (e.g., Jacob et al., 2000; Nweeia et al., 2012). The increasing availability of whole genome sequences, on the other hand, provides novel tools to untangle genomic signatures impacting organ reduction or loss (e.g. Zoonomia Consortium, 2020). A key question is thus to understand how genomic changes impact these processes. Among such signatures we find, more commonly than initially anticipated, gene loss episodes: such as in the morphological simplification of the urochordate *Oikopleura dioica*, the eye regression observed in cave-dwelling populations of the teleost *Astyanax mexicanus*, the loss of gastric glands in disparate vertebrate species or the loss of sebaceous glands in some mammalian lineages (e.g., Olson, 1999; Castro et al., 2014; Albalat & Cañestro, 2016; Cronk, 2009, Guijarro-Clarke et al. 2020; McGaugh et al., 2014; Springer et al. 2018; Lopes-Marques et. a 2019a; Themudo et al., 2020; Springer et al., 2021).

A remarkable example of inconsistent observations in the functionality *versus* vestigiality of an organ can be found in the anatomical observations of the pineal gland in mammals (Ralph, 1975). The pineal gland is a small endocrine organ present in the brain and playing a central role in the development of entrainment behaviors through the action of melatonin (circadian rhythmicity). From a physiological standpoint, melatonin synthesis occurs in a specialized cell type, the pinealocyte, through an enzymatic cascade involving the arylalkylamine *N*-acetyltransferase (*Aanat*) and *N*-acetylserotonin methyltransferase (*Asmt*) enzymes (Klein et al., 1997; Simonneaux & Ribelayga, 2003); subsequent signaling uses a set of high affinity receptors, *Mtnr1A* and *Mtnr1B*, involved in the response of the clock machinery to melatonin stimulation, leading to local and overt phase shifts (Figure 1) (Axelrod et al., 1964; Lewy et al., 1980; Reppart et al., 1996). Although anatomical studies clearly support a well-defined pineal gland in most mammals, in lineages such as cetaceans, mole rats and sirenians a true pineal gland seems to be absent; yet, some equivocal observations exist, ranging from complete absence to detectable presence of this gland in some species or in individuals within a species (Ralph, 1975; Ralph et al., 1985; Kim et al., 2011; Panin et al., 2012). Conflicting evidence reporting measurable levels of circulating melatonin (i.e. bottlenose dolphin) shed further doubt (Panin, 2012). Interestingly, gene loss signatures were identified in these lineages, supporting the loss-of-function of melatonin synthesis, a hallmark of pineal function, and/or signalling (Fang et al., 2014; Huelsmann et al., 2019; Lopes-Marques et al., 2019b), further demonstrating the power of genome analysis towards the clarification of organ function. The presence of a functional pineal gland is also contentious in Xenarthrans (armadillos, anteaters and sloths), a relatively understudied taxonomic group characterized by its intriguing nature (Figure 1; Oksche, 1965; Benítez et al., 1994; Superina & Loughry, 2015; Freitas et al., 2019) and representing one of the earliest-branching clades of placental mammals (Murphy et al., 2007; O’Leary et al., 2013). Xenarthrans are considered *imperfect* homeotherms, given their poor ability to adjust body temperature (Mc Nab, 1979; 1980; 1985). This inaptitude for thermal regulation, possibly related with their low metabolic rate and low energetic content diet, makes Xenarthrans’ activity patterns highly affected by air temperature, with potential effects in their circadian cycles (Chiarello, 1998; Giné et al., 2015; Maccarini et al., 2015; Di Blanco et al., 2017). While a recent report clearly identified pineal glands in the six-banded armadillo (*Euphractus sexcintus*), Linnaeus’s two-toed sloth (*Choloepus didactylus*), and in the southern tamandua (*Tamandua tetradactyla*), a distinct pineal was not found or was reported missing in species such as southern long-nosed armadillo (*Dasypus hybridus*), pale-throated sloth (*Bradypus tridactylus*), giant anteater (*Myrmecophaga tridactyla*) or big hairy armadillo (*Chaetophractus villosus*) (Benítez et al., 1994; Ferrari, 1998; Freitas 2019). However, in the nine-banded armadillo (*Dasypus novemcinctus*) inconsistent reports advocate for either the presence or absence of a genuine pineal gland (Harlow et al., 1981; Freitas et al., 2019). Also, variable concentrations of circulating serum melatonin during the 24 h day-night cycle have been detected in this species, raising the hypothesis of an extrapineal source for melatonin production (Figure 1; Harlow et al., 1981; 1982). With the emergence of various whole-genome sequences from Pilosa (sloths and anteaters) (e.g., Uliano-Silva et al., 2019) and Cingulata (armadillos) (e.g., Lindblad-Toh et al., 2011), Yin (et al., 2021) have recently reported the molecular erosion of *Aanat* in Xenarthra; yet, no attempt was made to expand this analysis to the full melatonin-related gene hub. Thus, we are now able to interrogate whether the gene repertoire of circadian rhythmicity is modified in this lineage and clarify the physiological status of the pineal gland within this group.

**Figure 1:**
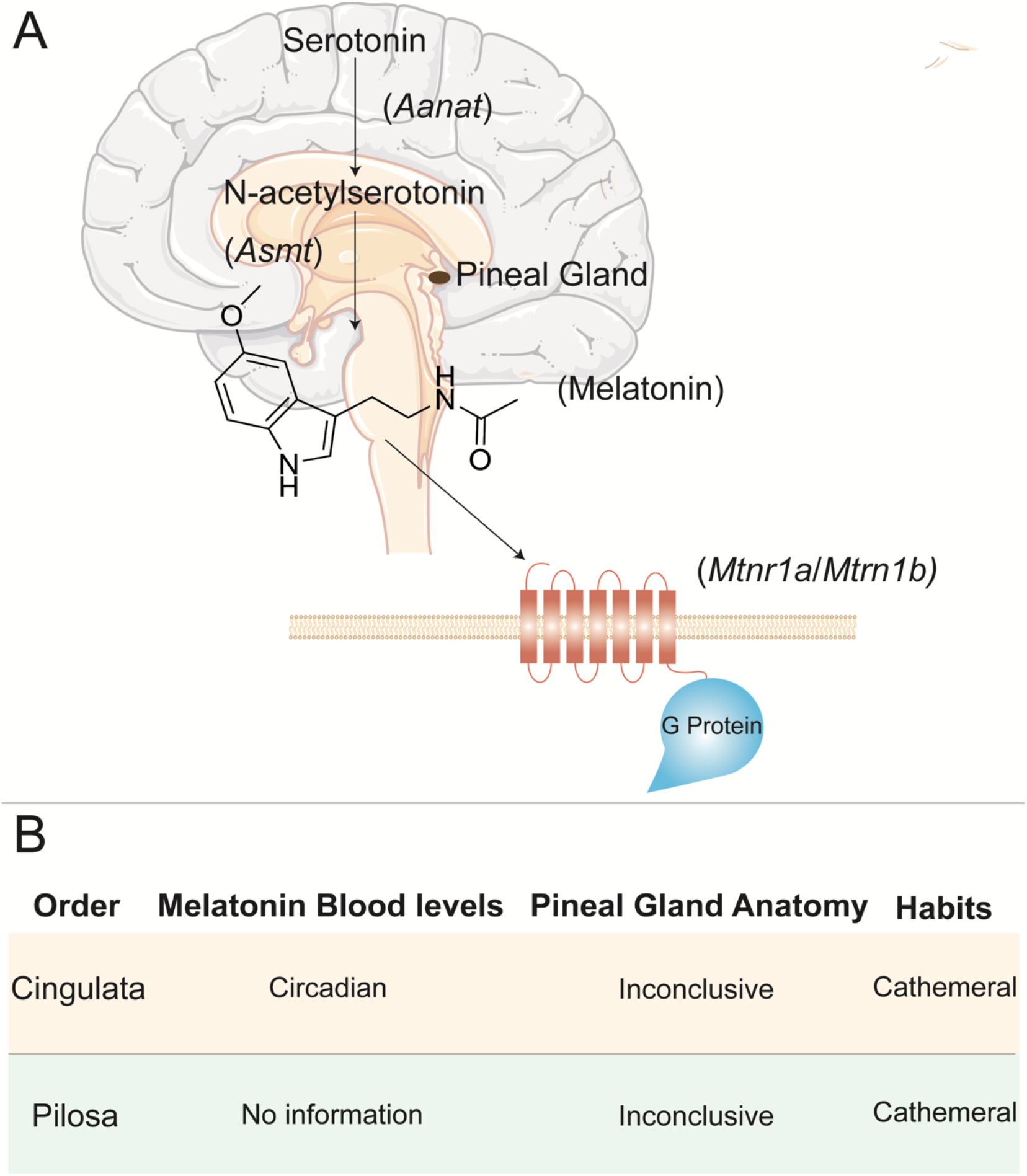
Melatonin synthesis and signaling. Melatonin, generally described as a phase marker of the circadian clock, is initially synthetized from tryptophan which is converted in serotonin (Pévet, 2002). The final steps of this synthetic pathway include a two-step metabolization of the intermediate serotonin into melatonin, a process catalyzed by *Aanat* and *Asmt* (Klein et al., 1997; Simonneaux & Ribelayga, 2003). In mammals, *Mtnr1a* and *Mtnr1b* receptors, are involved in the response of the clock machinery to melatonin stimulation (Reppart et al., 1996) (A). Summary of the available information regarding Xenarthran’s melatonin levels, pineal gland presence and habits (B). (Illustrations used elements from Servier Medical Art: https://smart.servier.com/)

## 2. Material and Methods

### 2.1 Sequence collection

To clarify the functional status of *Aanat, Asmt, Mtnr1a* and *Mtnr1b* in 8 Xenarthran species (Supplementary Data 1), the genomic *loci* were retrieved for gene annotation using three strategies (e.g. Alves et al., 2019; Lopes-Marques et al., 2019b): a) for species with annotated genes, the genomic sequence of the target gene (ranging from the upstream to the downstream flanking genes) was collected directly from NCBI; b) for species with annotated genomes but lacking the annotation of the target gene, the genomic region between two conserved flanking genes (downstream and upstream) was directly collected and c) for unannotated genomes, blastn searches were performed, using as query a set of three genes, including *Homo sapiens* (human) target gene coding sequence (CDS), as well as those of the flanking genes in the same species. From the blast results, the best matching genome scaffold corresponding to the consensus hit across those obtained per each query sequence was retrieved. When no consensual blast hit was obtained, all hits corresponding to the *H. sapiens* CDS query were inspected, the aligning regions submitted to a back-blast search against the nucleotide database of NCBI, with the matching genomic sequence corresponding to the gene of interest being the one selected (when existing). When several matchings were found, the best genomic scaffold (yielding the highest query coverage and identity value) was collected for annotation.

For the 156 non-Xenarthran mammals with annotated genomes (Supplementary Data 1), the first two strategies described above were adopted to obtain the genomic region corresponding to the target gene. In Dugong *(Dugong dugon)*, since no annotation is currently available, the genomic sequence containing the target gene was retrieved *via* blastn searches.

### 2.2 Gene Annotation and Mutational Validation

The open reading frames of the mammalian orthologues of *Aanat, Asmt, Mtnr1a* and *Mtnr1b* were investigated using Pseudo*Checker* (pseudochecker.ciimar.up.pt), an online platform suitable for gene inactivation inference (Alves et al., 2020). For this purpose, the human gene orthologue was used as a comparative coding sequence input (NCBI Accession ID regarding human *Aanat*: NM_001088.3; *Asmt*: NM_001171039.1; *Mtnr1a*: NM_005958.4; *Mtnr1b*: NM_005959.5) to deduce the coding status of a given candidate gene in each target species. By making use of PseudoIndex - a user assistant metric built into the Pseudo*Checker* pipeline that rapidly estimates the erosion condition of the tested genes – putative ORFs of the orthologous gene from each target species were assigned a discrete value from 0 to 5, with 0 suggesting a fully functional gene and 5 complete inactivation (Alves et al., 2020). When PseudoIndex was higher than 2, we proceeded to manual annotation and validation of possible disrupting mutations as previously described by Lopes-Marques et al. (2019a, 2019b, 2019c). Briefly, by using *H. sapiens* CDS for each target gene as reference, each exon was isolated and mapped to the genomic region of the candidate pseudogenes using Geneious Prime (2019.2.3) “map to reference” tool. The aligned regions were individually screened for ORF disrupting mutations (exon deletions, sequence frameshifts and premature stop codons) and identified mutations were annotated. Mutational validation was performed through retrieval of raw sequencing reads in (at least) two independent Sequence Read Archive (SRA) projects (when available).

### 2.3 RNA-seq analysis

Transcriptomic analysis was performed as previously described by Lopes-Marques (et al., 2019c). Succinctly, RNA-seq datasets of multiple tissues were obtained from SRA projects to inspect the functional condition of each target gene in Xenarthran species (when available) and Human (*H. sapiens*) (Supplementary Data 2). Transcriptomic reads recovered through blastn, were mapped to corresponding references genomes using the “map to reference” tool from Geneious Prime (2019.2.3) and manually removed if presenting poor alignment. Finally, reads were classified as spliced reads (spanning over two exons), exon-intron reads or exonic reads depending on the genomic region they mapped.

## 3. Results

### 3.1 Erosion of melatonin-related genes in Xenarthra

To infer the coding state of the melatonin synthesis genes in armadillos, anteaters and sloths, we compared the genomic regions containing *Aanat* and *Asmt* in *H. sapiens* to the full genomes of 8 eight Xenarthran species and *L. africana* (Supplementary Data 1). Analysis using Pseudo*Checker* (Alves et al., 2020), showed that all analysed species presented a PseudoIndex equal to 5 (Supplementary Data 3) - thus suggesting that the ORF of *Aanat* and *Asmt* includes inactivating mutations. Subsequent manual annotation of all collected Xenarthran genomic sequences revealed *Aanat* and *Asmt* gene erosion across all analysed species (Supplementary Data 4). Regarding *Aanat*, in agreement with Yin (et al., 2021), we found numerous ORF-disrupting mutations, including a conserved 1-nucleotide deletion in exon 1 in Armadillos and a conserved 2-nucleotide deletion in exon 3 in Sloths (Figure 2; Supplementary Data 4). On the other hand, although Yin (et al., 2021) were not able to recover the genomic sequence containing the *Aanat* CDS in Anteaters, we uncovered, among other disruptive mutations, a conserved in-frame premature stop codon in exon 2. The identified mutations were next validated by searching at least one ORF disruptive mutation per species in the corresponding SRAs; reads corroborating the identified mutations were systematically found (Supplementary Data 5). The analysis of *Asmt* in Xenarthra also revealed variable disruption patterns across Xenarthra. In cingulatans, we found similar mutational events, however not conserved within members of this group. Specifically, in *D. novemcinctus* exons 1 to 4 and 7 were not found, possibly due to poor genome coverage or complete exon deletion (Figure 2). Moreover, in southern three-banded armadillo (*Tolypeutes matacus*) several insertions/deletions (indels) have been identified in exon 6, contrasting with *D. novemcinctus* where a validated in-frame premature stop codon in the same exon was detected (Figure 2; Supplementary Data 4 and 6). In Vermilingua (anteaters), we were only able to recover exons 4 and 5 in the Giant anteater (*Myrmecophaga tridactyla*) which provide a range of mutations with predicted disruptive effects (Figure 2; Supplementary Data 4). For Folivora (sloths), across several identified ORF-disrupting mutations, a trans-species conserved 4-nucleotide insertion in exon 6 was revealed and further validated by SRA searches (Supplementary Data 4 and 6). RNA-Seq analysis in Linnaeus’s two-toed sloth (*Choloepus didactylus*) *Aanat* yielded a high proportion of exon-intron reads versus spliced reads, in clear contrast with the pattern found in *H. sapiens* (Supplementary Data 7). In the case of *Asmt*, no transcriptomic reads were recovered for *C. didactylus*. Similarly, SRA transcriptome searches were unable to retrieve reads of *D. novemcinctus* for both genes.

**Figure 2:**
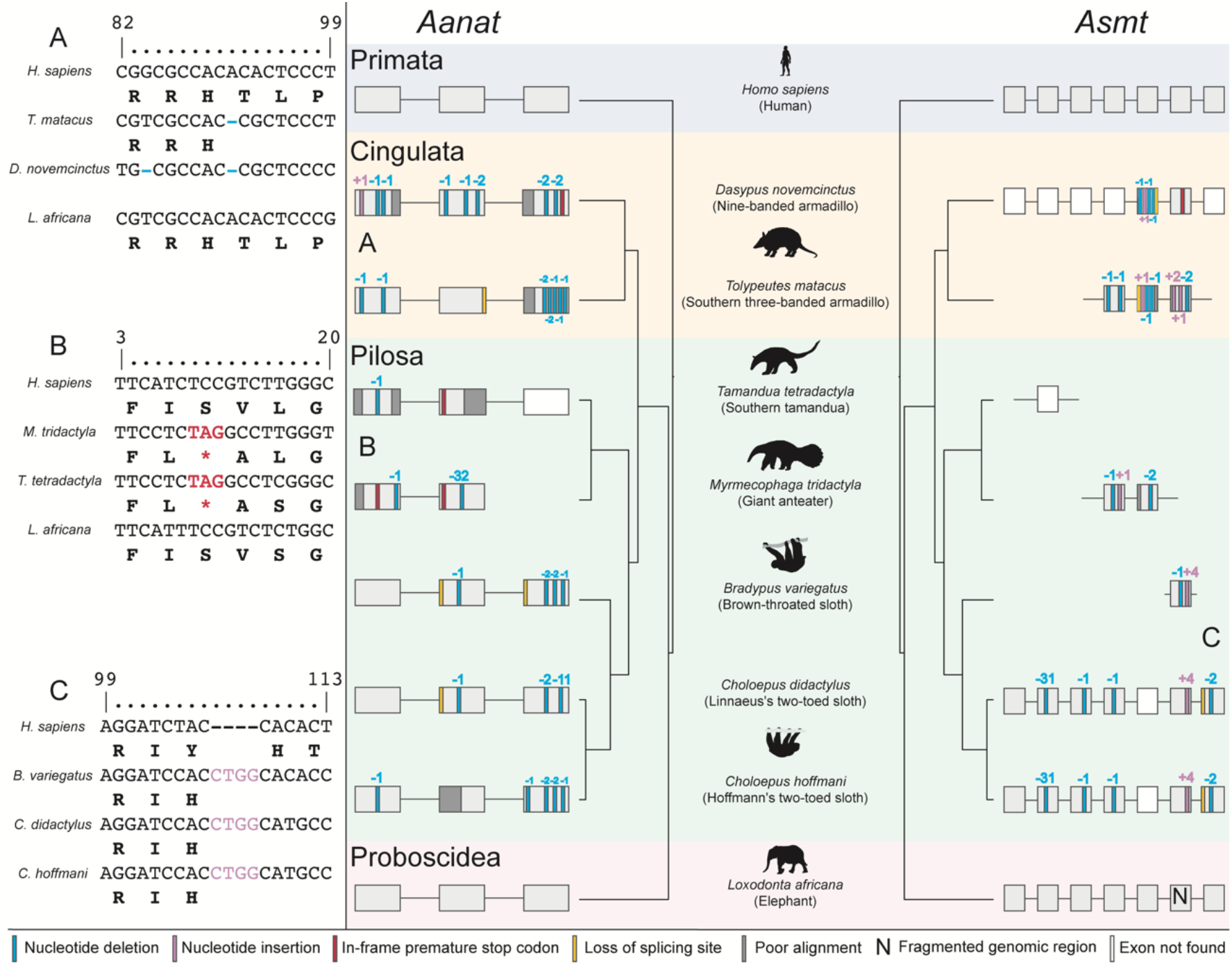
Schematic representation of the identified *Aanat* and *Asmt* genes ORF abolishing mutations in Xenarthra orders (Cingulata and Pilosa). Phylogenetic trees were calculated in www.timetree.org; last accessed March 13, 2021 using species list. Silhouettes were sourced from Phylopic (http://phylopic.org). Sequence alignments of the identified conserved disruptive mutations in both *Aanat* and *Asmt* genes of Xenarthra.

We next examined the genes *Mtnr1a* and *Mtnr1b*, that encode G-protein coupled receptors responsible for melatonin signalling. In *D. novemcinctus*, the *Mtnr1a* coding status could not be accessed likely due to fragmentation of the respective genomic region (presence of sequencing gaps (Ns)). On the other hand, for both species comprising the two-toed sloth group (*Choloepus* sp.), we were able to identify a validated 8-nucleotide deletion and a 20-nucleotide deletion in exon 2 together with a 4-nucle0tide insertion in exon 1 (Figure 3; Supplementary Data 4 and 8). Curiously, in elephant (*Loxodonta africana*) exon 2 was not found despite the completeness of the assembly in the *Mtnr1a* region. The analysis of *Mtnr1b* CDS in *T. matacus* and Screaming hairy armadillo (*Chaetophractus vellerosus*) uncovered several inactivating mutations, including a conserved premature stop codon that truncates exon 2 (Figure 3). Pilosa species (sloths and anteaters) *Mtnr1b* gene annotation revealed the presence of several ORF disrupting mutations, of note a single transversal mutation present in all analysed species, namely a 2-nucleotide deletion in exon 2 (Figure 3; Supplementary Data 4). This mutation was investigated and validated in both *Choloepus* species (Supplementary Data 9). Searches for transcriptomic evidence for *Mtnr1a* and *Mtnr1b* in *C. didactylus* retrieved a low number of reads, mostly corresponding to immaturely mRNA (Supplementary Data 7).

**Figure 3:**
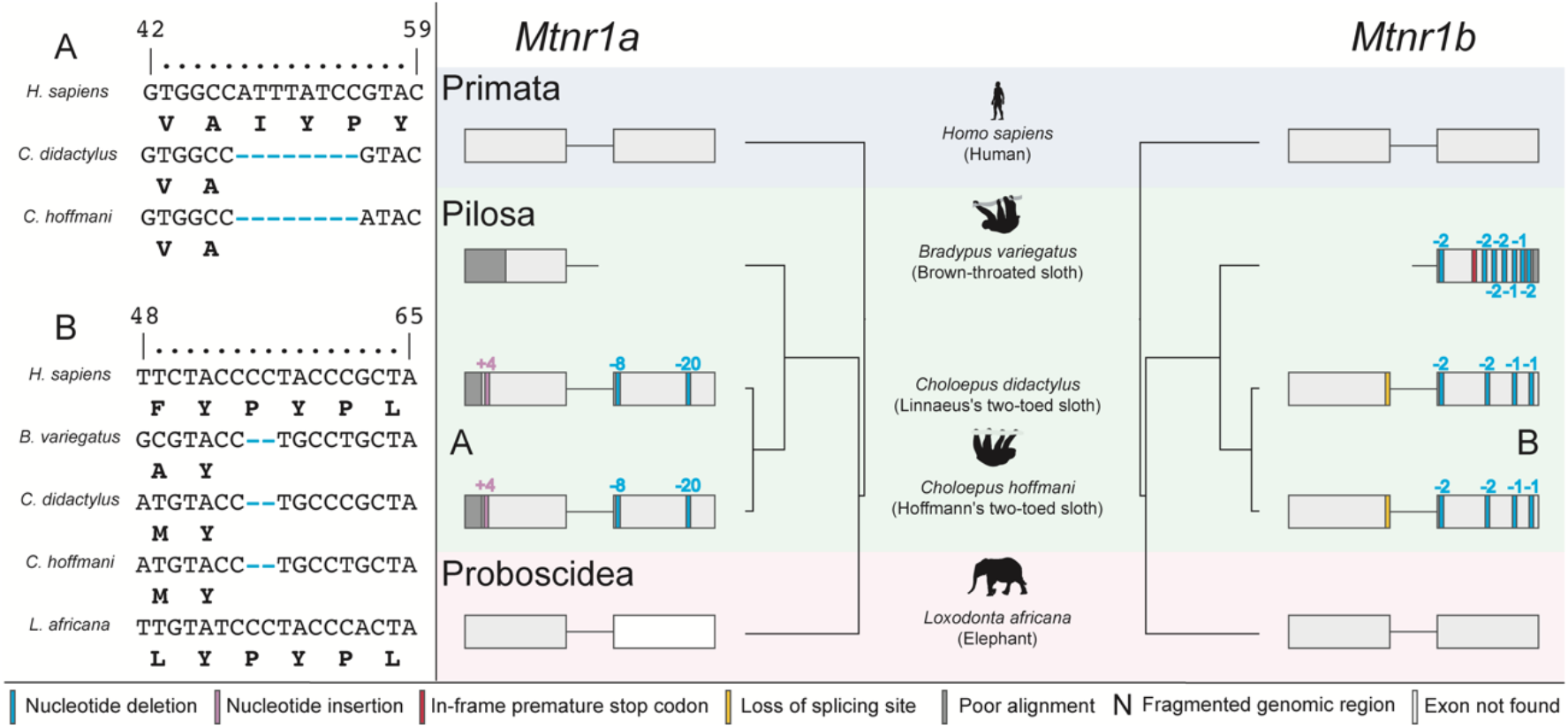
Schematic representation of the identified *Mtnr1a* and *Mtnr1b* genes ORF abolishing mutations in Xenarthra orders (Cingulata and Pilosa). Phylogenetic trees were calculated in www.timetree.org; last accessed March 13, 2021 using species list. Silhouettes were sourced from Phylopic (http://phylopic.org). Sequence alignments of the identified conserved disruptive mutations in both *Mtnr1a* and *Mtnr1b* genes of Xenarthra.

### 3.2 Melatonin-related genes are inactivated in other non-xenarthran mammals

We next expanded our analysis to non-Xenarthran mammal genomes (157), to address the coding status of *Aanat, Asmt, Mtnr1a* and *Mtnr1b* (Supplementary Data 1). Sequence search and analysis for *Aanat* returned a total of 11 species with no annotation of a *Aanat*-like sequence: *Bison bison bison* (American bison), *Bos indicus* (Zebu), *Bos mutus* (Wild yak), *Bubalus bubalis* (Water buffalo), *Camelus ferus* (Wild bactrian camel), *Odocoileus virginianus texanus* (White-tailed deer), *Pantholops hodgsonii* (Tibetan antelope), *Sus scrofa* (Wild boar), *Myotis davidii* (David’s myotis) and *Myotis lucifugus* (little brown bat). For the latter, we were not able to retrieve the genomic locus containing the target gene, given that both upstream and downstream flanking genes are also not annotated. Analysis using Pseudo*Checker* (Alves et al., 2020), showed that 32 species non-Xenarthran mammals presented a PseudoIndex higher than 2 (Supplementary Data 3). From these species, members of Cetacea (cetaceans) and Pholidota (pangolins) presented among their members, a conserved (and validated) in-frame premature stop codon in Exon 1 (Supplementary Data 4 and 10). Moreover, we also found ORF-disrupting mutations in Exon 1 of velvety free-tailed bat (*Molossus molossus*), Kuhl’s pipistrelle (*Pipistrellus kuhlii*) and Sunda flying lemur (*Galeopterus variegatus*) with the latter being validated through SRA genomic reads (Supplementary Data 4 and 10). In *D. dugon*, several disruptive mutations were identified, such as an eight-nucleotide insertion in exon 2 and the presence of a stop codon in exon 3 (Supplementary Data 4 and 10).

Regarding *Asmt*, in 17 species the genomic fragments containing *Asmt*-like nucleotide sequences were not recovered given the lack of annotation for both target and flanking genes (Supplementary Data 3). For this gene, 74 species displayed a PseudoIndex higher than 2 (Supplementary Data 3), the majority due to fragmentation of the genomic region (presence of Ns), true absence of exons, poor alignment identity or incompleteness of the scaffold in the *Asmt* genomic region (Supplementary Data 4). Gene lesion events were found and validated mostly in Cetaceans, *G. variegatus, Manis sp*. (pangolins) and some Rodentia, with the latter showing poor alignment identity with the reference (Figure 4; Supplementary Data 4 and 11). Other examples of species presenting disruptive mutations but with no SRA validation (given that no genomic independent SRAs projects are available) include Brandt’s bat (*Myotis brandtii*), the Prairie vole (*Microtus ochrogaster*) with 2-nucleotide deletion in exon 1 and the Nancy Ma’s night monkey (*Aotus nancymaae*) with single nucleotide deletions in exon 2 (Supplementary Data 4). In the case of *D. dugon*, no ORF-disrupting mutations were found for *Asmt*, however, not all the exons were recovered due to incompleteness of the scaffold (Supplementary Data 4).

**Figure 4:**
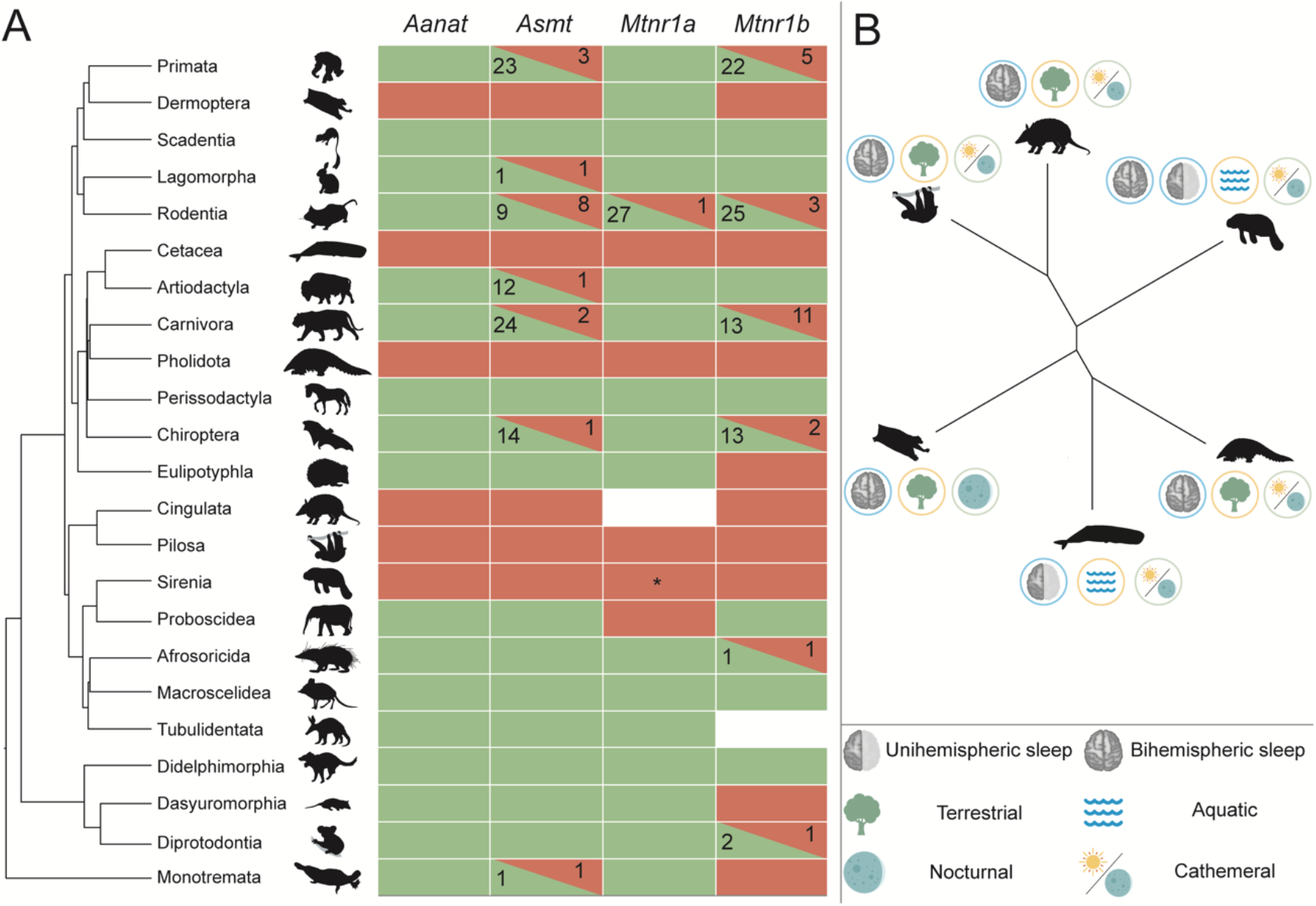
Mutational landscape of melatonin synthesis and signalling genes along the mammalian tree. For each gene, we represented in green the orders where no ORF-disrupting mutations (frameshift mutations, in-frame premature stop codons, loss of canonical splicing site or exon deletions) were found across all members. On the other hand, orders where all members presented ORF-disrupting mutations are highlighted in red. In orders (and in genes) with no consensual disruption pattern, number of species presenting a coding/non-coding sequence were depicted respectively. Species where no SRA validation was possible, were not included in this figure. * indicates the presence of contradictory reports in *Mtnr1a* for *Trichechus manatus latirostris*. Phylogenetic relationships were adapted from Vazquez et al. (2018). (A) Summary characterization of mammalian lineages presenting complete molecular erosion of melatonin synthesis and signalling genes, regarding their sleep type, habitat and lifestyle. (B)

We next expanded our search to understand whether melatonin signaling genes would be compromised in non-Xenarthran mammals. For *Mtnr1a*, a total of 44 species exhibited a PseudoIndex higher than 2 (Supplementary Data 3), yet manual curation revealed ORF-disrupting mutations only in pangolins (validated through SRA Projects; Supplementary Data 12) (Huelsmann et al., 2019), Hawaian monk seal (*Neomonachus schauinslandi*) and *D. dugon* (Supplementary Data 4). In cetaceans a different scenario emerged, with several exons completely absent (Supplementary Data 4) (Huelsmann et al., 2019; Lopes-Marques et al., 2019b).

In *Mtnr1b*, a total of 69 non-Xenarthran mammals displayed a PseudoIndex higher than 2 (Supplementary Data 3). However, and contrary to the pattern found for *Mtnr1a*, annotation of the collected sequences revealed *Mtnr1b* gene erosion across multiple species, mostly affecting the Carnivora and Cetaceans but also Pholidota, Sirenia and some Primates (Figure 4; Supplementary Data 4). Examples of conserved inactivating mutations were found in bears (*Ursus sp*.) with an in-frame premature stop codon in exon 1, weasels (*Mustela sp*.) sharing several indels in exon 2 and pangolins with a single nucleotide deletion and an in-frame premature stop codon in exon 1 (Supplementary Data 4 and 13). Other species with ORF-disruptive mutations include *Nannospalax galili* (northern Israeli blind subterranean mole rat), exhibiting a single nucleotide deletion in exon 1, *D. dugon* with a conserved fourteen-nucleotide deletion in exon 2 or *G. variegatus* with an indel also in exon 1 (Supplementary Data 4 and 13). Detailed characterization of each target gene in mammals is available in Supplementary Data 4 and the minutiae of SRA validation can be found in Supplementary Data 10, 11, 12 and 13.

## 4. Discussion

Here, we set out to investigate how evolutionary genomic signatures might untangle the physiological status of controversial vestigial structures, using the pineal gland as a case study (Pévet, 2002). For this, we addressed the evolution of a melatonin-related gene hub, encompassing melatonin synthesis and signaling genes, in Xenarthra and other mammals. Our results strongly suggest a complete landscape of gene loss in Xenarthra, which further reinforce reports suggesting the lack of a pineal gland in several members of this superorder (Quay, 1965; Harlow et al., 1981; Benítez et al., 1994, Ferrari 1998; Freitas 2019). On the other hand, in species in which a pineal gland was described (e.g., Freitas et al., 2019), the present data suggests that, despite the anatomical observations, the canonical pineal gland physiology leading to melatonin secretion is likely disrupted. Nevertheless, similarly to what was described for *Tursiops truncatus* (bottlenose dolphin) (Panin et al., 2012), previous radioimmunoassay methods have reported the presence of melatonin circulating in *D. novemcinctus* (Harlow et al., 1981), implying either the existence of independent pathways for melatonin synthesis and signaling (Slominski et al., 2003; Tan et al., 2016) or possible acquisition of melatonin from food sources (Tan et al., 2010).

This strong genomic signature leading to the anatomical and/or physiological atrophy of this endocrine gland might be viewed as an adaptive solution to overcome physiological limitations. Described as cathemeral (irregular daily activity pattern) and heterothermic species (Eisenberg & Redford, 1999) with limited capacity to regulate their body temperature, Xenarthrans’ movements are heavily influenced by air temperature (Greegor, 1985; Camilo-Alves & Mourão, 2006; Giné et al., 2015; Attias et al., 2018). Thus, to reduce such energetic costs, Xenarthrans may have suffered reductive episodes, allowing behavioral strategies to overcome unfavorable environmental conditions and mitigate thermal limitations (Yin et al., 2021).

Accordingly, convergent disruptive patterns with lineages also presenting labile body temperature and suggestive bizarre sleeping patterns (Pholidota; Mc Nab, 1984; Heath & Hammel, 1986; Weber et al., 1986; Imam et al., 2018; Yin et al., 2021) or living in environments with specific thermal constraints (Cetacea and *Trichechus manatus latirostris* (Florida manatee); Huelsmann et al., 2019; Lopes-Marques et al., 2019b; Yin et al., 2021), make it plausible to hypothesize that, in these species, inactivation of melatonin-related genes can be related with changes in their circadian rhythmicity. In addition, pseudogenization of these genes possibly paralleled loss of other circadian rhythm related genes, namely Cortistatin gene, that encodes a pleiotropic neuropeptide with an important role in sleep physiology (Valente et al., 2021). Given that the evolution of melatonin-related genes should be directly linked with pineal gland function, by inferring their coding status we were able to deduce if the organ constitutes an evolutionary vestige, despite the conflicting anatomical reports. More importantly, the present study provides a clear case-study on how genomic data can be used to disentangle whether a specific organ constitutes a functional or vestigial structure (Hiller et al., 2012).

## 5. Conclusion

To date, no unequivocal inferences on the functional status of pineal gland across mammals were provided, with anatomical observations in several species from different clades presenting conflicting conclusions. However, by making use of genomic data, our results provide solid evidence for pineal gland vestigiality not only in Xenarthra, but also in other mammalian lineages. Thus, we argue that analysis of genomic changes might constitute a powerful approach to gain insights into the vestigiality of specific organs.

## Author’s contributions

**Raul Valente:** Data curation, Formal analysis, Investigation, Methodology, Visualization, Writing - original draft **Filipe Alves:** Writing - review & editing **Isabel Sousa-Pinto:** Writing - review & editing **Raquel Ruivo:** Conceptualization, Methodology, Validation, Writing - review & editing **Luís Filipe Costa de Castro:** Conceptualization, Methodology, Validation, Supervision, Project administration, Resources, Writing - review & editing

## Acknowledgements

We acknowledge the various genome consortiums for sequencing and assembling the genomes.

